# Interpreting coronary artery disease risk through gene-environment interactions in gene regulation

**DOI:** 10.1101/475483

**Authors:** Anthony S Findley, Allison L Richards, Cristiano Petrini, Adnan Alazizi, Elizabeth Doman, Alexander G Shanku, Omar Davis, Nancy Hauff, Yoram Sorokin, Xiaoquan Wen, Roger Pique-Regi, Francesca Luca

## Abstract

GWAS and eQTL studies identified thousands of genetic variants associated with complex traits and gene expression. Despite the important role of environmental exposures in complex traits, only a limited number of environmental factors are measured in these studies. Measuring molecular phenotypes in tightly controlled cellular environments provides a more tractable setting to study gene-environment interactions in the absence of other confounding variables.

We performed RNA-seq and ATAC-seq in endothelial cells exposed to retinoic acid, dexamethasone, caffeine, and selenium to model genetic and environmental effects on gene regulation in the vascular endothelium, a common site of pathology in cardiovascular disease. We found that genes near regions of differentially accessible chromatin were more likely to be differentially expressed (OR = [3.41, 6.52], p < 10^−16^). Furthermore, we confirmed that environment-specific changes in transcription factor binding are a key mechanism for cellular response to environmental stimuli. SNPs in these transcription response factor footprints for dexamethasone, caffeine, and retinoic acid were enriched in GTEx eQTLs from artery tissues indicating that these environmental conditions are latently present in GTEx samples. Additionally, SNPs in footprints for response factors in caffeine are enriched in colocalized eQTLs for coronary artery disease (CAD), suggesting a role for caffeine in CAD risk. Interestingly, each treatment may amplify or buffer genetic risk for CAD, depending on the particular SNP considered.

## Introduction

The vascular endothelium refers to the single layer of cells lining blood vessels which form an interface between the blood and the rest of the body. The endothelium plays an important role in coagulation, thrombosis, leukocyte extravasation, regulation of vascular tone, and angiogenesis [42]. Endothelial dysfunction has been implicated in many pathological processes, including atherosclerosis, hypertension, tumor angiogenesis, wound healing, and preeclampsia [21]. For many of these conditions, genome-wide association studies (GWAS) have been instrumental in identifying a large number of genetic variants associated with disease (e.g. [37]). However, the molecular mechanisms linking each variant to the disease phenotype are largely unknown. This is because risk variants tend to fall in non-coding regions and affect regulatory mechanisms that are not yet well characterized.

DNase-seq and ATAC-seq [5, 6] are experimental approaches developed to profile the chromatin landscape and can be used to identify active regulatory elements. Transcription factors bound to chromatin leave characteristic ‘footprints’, allowing for the identification of many bound transcription factors in a single experiment [41]. ATAC-seq offers the advantage of requiring a much smaller number of cells, allowing for increased multiplexing of experiments in small volumes. This enables study of cell lines with differing genetic backgrounds across environmental perturbations, such as the addition of a drug or hormone. Current existing annotations for genetic variants only capture regulatory mechanisms in one arbitrary environment and thus it is necessary to annotate regulatory regions across multiple conditions. Combining chromatin accessibility data with measures of gene expression, such as RNA-seq, provides insights into the regulatory code governing gene expression. By comparing RNA- and ATAC-seq in the presence/absence of perturbations, it becomes possible to comprehensively study the cellular response and link changes in chromatin accessibility and transcription factor binding to changes in gene expression [2, 19, 36, 39]. The molecular basis of cellular response can be identified by leveraging genetic differences within and across individuals to pinpoint underlying regulatory elements. One such approach using genetic variation is quantitative trait loci (QTL) mapping, whereby genetic variants can be linked to differences in gene expression (eQTL) (e.g. [1, 49] or chromatin accessibility (caQTL) [9], among others). Results from these methods, which can be used individually or as complements to one another, have been integrated with GWAS findings to suggest a model in which genetic variants disrupt the chromatin architecture, leading to differences in gene expression which affect complex traits[18, 22, 25, 54].

However, the effect of a genetic variant on a molecular pathway, and ultimately on the disease condition, may be modulated by environmental factors [27, 28, 35]. Known environmental risk factors for endothelial dysfunction include diabetes, smoking, obesity, high cholesterol, and hypertension [23]. Dexamethasone, retinoic acid, caffeine, and selenium are four treatments that have been associated with CVD and/or endothelial dysfunction[17, 40, 50, 52] and that produce strong transcriptional responses in endothelial cells[35]. Dexamethasone is a potent glucocorticoid that serves as a proxy for stress [46] and excess may promote calcification within arteriosclerotic lesions [56]. Retinoic acid plays a crucial role in the development of the cardiovascular system, and vascular endothelial cells are exposed to high concentrations of retinoic acid and express retinoid receptors [45]. Caffeine, the most widely consumed stimulant worldwide, promotes vasodilation in the endothelium and has been the subject of numerous studies regarding its association with CVD with conflicting results [12, 51]. Selenium is a component of selenoproteins which act as antioxidants, and selenium has been used as a supplement to increase HDL cholesterol [43].

Recently, the heterogeneity of the contexts represented in complex datasets has been used as proxy for varying environmental conditions [28, 55] to identify GxE effects. The advantage of these approaches resides in the possibility of exploiting pre-existing large-scale eQTL studies that profile gene expression in tissues from a heterogeneous sample of individuals; however the differences in environmental conditions between individuals is not explicitly ascertained. To characterize genetic and environmental risk factors for cardiovascular disease, it is critical to investigate gene regulation in a relevant cell type under controlled treatment conditions. Here we developed annotations of genetic variants in key regulatory elements for gene expression response to inform the role of gene-environment interactions (GxE) in coronary artery disease.

## Results

### Exposure of endothelial cells to environmental perturbations leads to changes in gene expression and chromatin accessibility

To determine the impact of genetic variation on endothelial cell response to environmental perturbations, we treated human umbilical vein endothelial cells (HUVECs) from 17 unrelated, healthy donors with 4 compounds (dexamethasone, retinoic acid, caffeine, and selenium) and 2 vehicle controls. These treatments have been associated with cardiovascular disease and/or endothelial dysfunction [17, 40, 50, 52] and have been shown in a previous study [35] to produce strong transcriptional responses in HUVECs. We then assessed changes in gene expression with RNA-seq in all samples and chromatin accessibility with ATAC-seq in four samples at six hours following treatment (Figure 1A).

**Figure 1:**
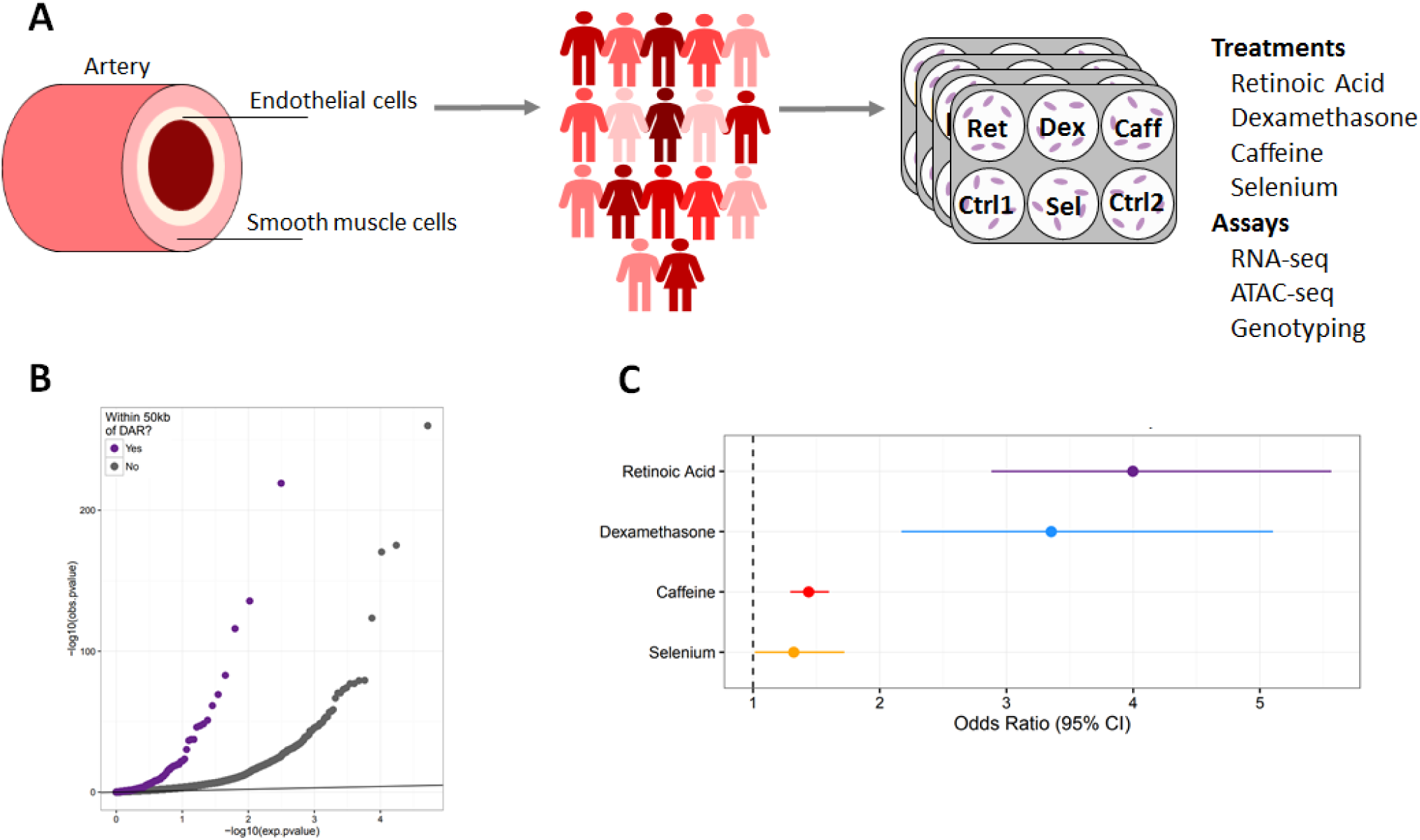
Changes in gene expression and chromatin accessibility in HUVECs treated with dexamethasone, retinoic acid, caffeine, and selenium. **A**Diagram of artery layers and study design. HUVECs were treated separately with 4 compounds and vehicle controls prior to RNA- and ATAC-seq (Methods). **B** QQ-plot of *p*-values from DESeq2 analysis of gene expression in response to retinoic acid. The purple dots represent genes within 50kb of a DAR, while the gray dots represent genes not within 50kb of a DAR. **C** Enrichment of DE genes within 50kb of a DAR. Odds ratio was estimated using Fisher’s exact test, and for all treatments *p <* 2.2 × 10^−^16.

We used DESeq2 [30] to identify differentially expressed genes (DE genes) and differentially accessible regions (DARs) in chromatin following the environmental perturbations. Each treatment caused significant transcriptional responses, with greater than 2500 differentially expressed genes in each environment (Benjamini-Hochberg (BH) FDR < 10%, log2 FC *>* 0.25) (Table 1). By gene ontology analysis, we identified an enrichment of differentially expressed genes implicated in biological processes relevant for endothelial function. For example, response to wounding is enriched in genes downregulated in response to dexamethasone, and regulation of blood vessel diameter is enriched in genes downregulated in response to retinoic acid (Supplementary Table S1).

**Table 1:** Number of differentially expressed genes and differentially accessible regions

| Treatment | DE Genes | DARs |
| --- | --- | --- |
| Retinoic Acid | 4,874 | 222 |
| Dexamethasone | 2,879 | 133 |
| Caffeine | 5,790 | 1,038 |
| Selenium | 10,329 | 196 |

Across the four samples for which ATAC-seq was obtained, we identified between 133 and 1038 DARs in a given treatment condition (Table 1). The smaller number of DARs, compared to DE genes, could be due to the timescale at which we measured response, the increased stochasticity of chromatin opening and closing relative to gene expression, and/or insufficient statistical power. We used chromHMM [16] chromatin states derived from ENCODE HUVECs histone marks [15] to annotate DARs. We found an enrichment of DARs in strong enhancer regions in dexamethasone, retinoic acid, and selenium (OR = 5.31, 3.89, and 3.51, respectively. p < 2.2 × 10^−16^ for all). However, caffeine DARs are only slightly enriched in strong enhancers (OR = 1.15, p = 0.018). Instead, DARs in caffeine are most enriched for active promoters (OR = 2.5, p < 2.2 × 10^−16^). This difference between treatments could indicate that different type of factors are involved in each response.

To investigate the relationship between differential chromatin accessibility and transcriptional response, we calculated the enrichment of DE genes within 50kb of a DAR. DE genes in all treatments were significantly enriched near DARs identified in the same treatment. The greatest enrichments were found for dexamethasone and retinoic acid with odds ratios of 3.35 and 3.99, respectively (Figure 1B,C and Supplementary Figure S1; Fisher’s exact test; p < 5.6 × 10^−8^). The significant enrichment for DE genes among DARs in dexamethasone and retinoic acid could be due to their mechanism of action, specifically that both treatments are known to work largely through specific nuclear receptors that bind the DNA and influence gene expression directly. Overall these results suggest that changes in chromatin accessibility are an important mechanism through which cells regulate gene expression in response to environmental perturbation.

### Specific transcription factors regulate response to treatment

In addition to informing on open chromatin, ATAC-seq data can be analyzed to characterize the transcription factor landscape governing response to treatment. This is particularly important given that the vast majority of genetic variants found to influence complex traits act through gene regulatory regions. In order to identify transcription factors controlling response to each treatment, we used CENTIPEDE [41] to perform footprinting analysis on the ATAC-seq data. Across all conditions, we identified a total of 882 active transcription factors, defined as those with motifs that have CENTIPEDE footprints and are associated with open chromatin (Methods, Supplementary Table S6, see examples in Figure 2A). We hypothesized that transcription factors which are important in the regulation of a particular treatment will be more likely to be found in DARs. The factors with the greatest enrichments in DARs are shown in Figure 2B. We refer to these enriched factors as response factors. We identified 724 factors with footprints in regions that become more accessible after treatment and 259 factors in regions that are less accessible after treatment (Table 2). For well-studied treatments, like retinoic acid and dexamethasone, our approach identified transcription factors known to be central to the regulation of gene expression in response to those treatments. For example, the top retinoic acid response factors were the retinoic acid receptor along with two interferon regulatory factors (IRF), all of which has previously been shown to be influenced by retinoic acid[4, 8, 31] (Figure 2A). For dexamethasone, the androgen receptor (AR), glucocorticoid receptor (GR), and progesterone receptor (PR) have similar binding sites and likely all represent sites of GR binding[47, 48]. While for these treatments we have prior information on the major mechanisms that regulate the cellular response, for caffeine and selenium less is known, and our data can be used to identify the relevant factors (Supplementary Table S3). In response to selenium treatment the top enriched factors are in regions that become less accessible after treatment. Caffeine has been shown to induce apoptosis in HUVECs [29, 32], and one of the top caffeine response factors is Sp1, which has been implicated in apoptotic pathways [10, 24].

**Table 2:** Number of response factors enriched in opening and closing chromatin and SNPs in response factors footprints

| Treatment | Opening Chromatin | Closing Chromatin | Annotated SNPs |
| --- | --- | --- | --- |
| Retinoic Acid | 159 | 1 | 956,277 |
| Dexamethasone | 77 | 2 | 662,391 |
| Caffeine | 488 | 131 | 2,376,297 |
| Selenium | 0 | 125 | 924,329 |

**Figure 2:**
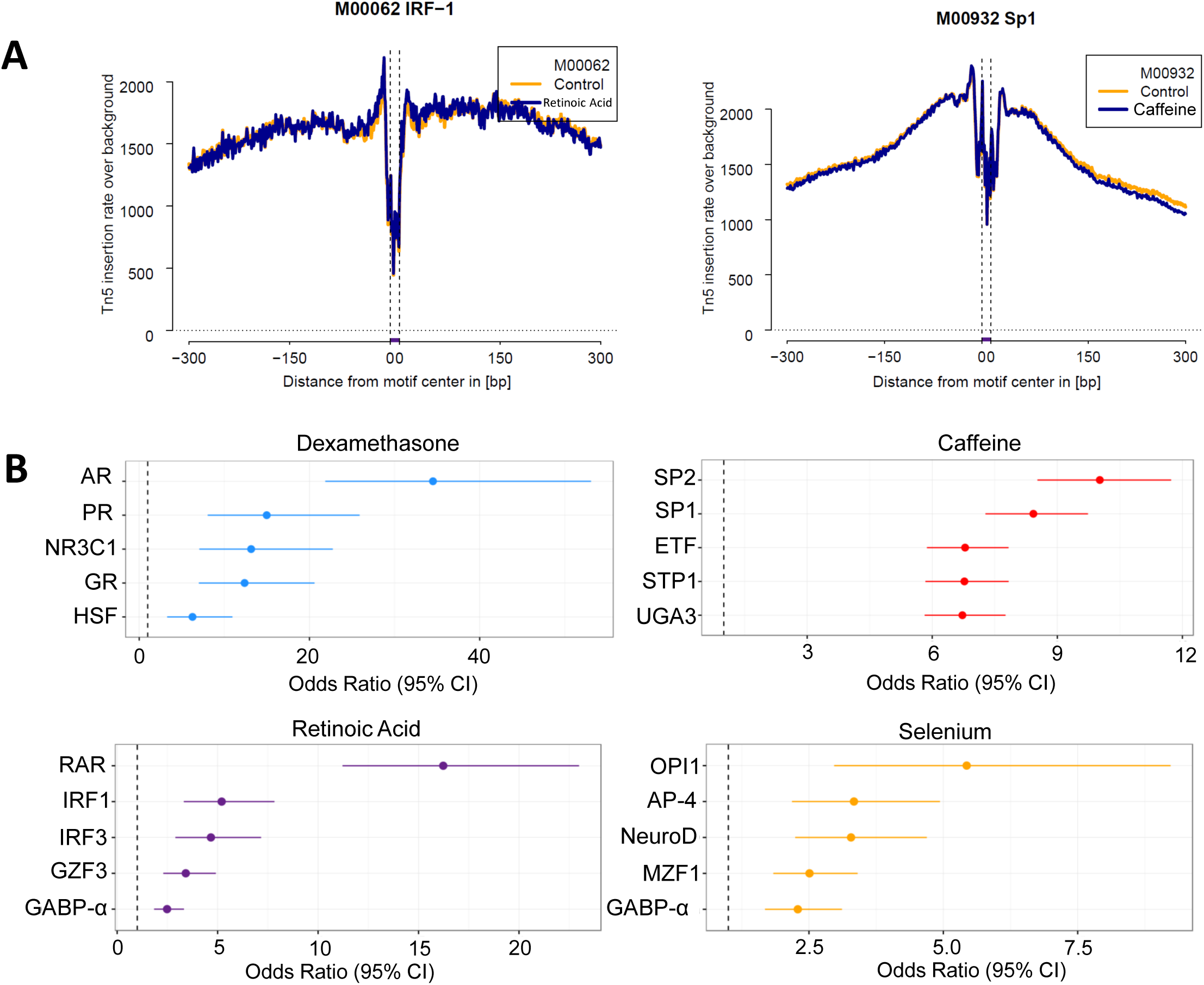
Identification of response factors. **A** Footprints of IRF-1 and Sp1 from ATAC-seq data in retinoic acid and caffeine treated cells, respectively. **B** Response factors, top most enriched transcription factor footprints in DARs in each condition. For selenium, the top factors are enriched in regions of closing chromatin. For all other treatments, the top factors are enriched in regions of opening chromatin.

### Identifying latent environmental factors in large scale eQTL datasets from vascular tissues

Large-scale efforts to identify expression quantitative trait loci (eQTLs), such as GTEx [1], have discovered genetic regulators of gene expression in many different tissues, including three types of artery. However, GTEx samples likely represent a composite of different environmental exposures. Here we propose that latent, unobserved environmental factors contribute to eQTL effects detected in GTEx and that some of these eQTLs may represent cases of significant interaction effects in specific environmental contexts. To indirectly test this hypothesis, we considered that interactions with latent environments represented by our four *in vitro* treatments (retinoic acid, caffeine, selenium and dexamethasone) would be enriched in response factor binding sites. We used centiSNPs [35], an extension of the CENTIPEDE framework, to annotate SNPs in response factor footprints, as these SNPs are most likely to disrupt transcription factor binding and modify the transcriptional response. We found that out of 4,422,115 SNPs in footprints for active factors, 76% are in footprints for response factors for at least one of the treatments. We then estimated enrichments for our treatment-specific annotations in eQTLs from the artery samples in GTEx. Using the EM-DAP1 algorithm implemented in the software package TORUS [53], we found enrichment of eQTLs in footprints for dexamethasone, retinoic acid, and caffeine response factors, but not for selenium (Figure 3A). These results suggest that exposure to retinoic acid, caffeine, and dexamethasone (or a closely related factor; e.g. cortisol elevated by stress) are represented as latent environments in the GTEx samples and can modulate eQTL effects.

**Figure 3:**
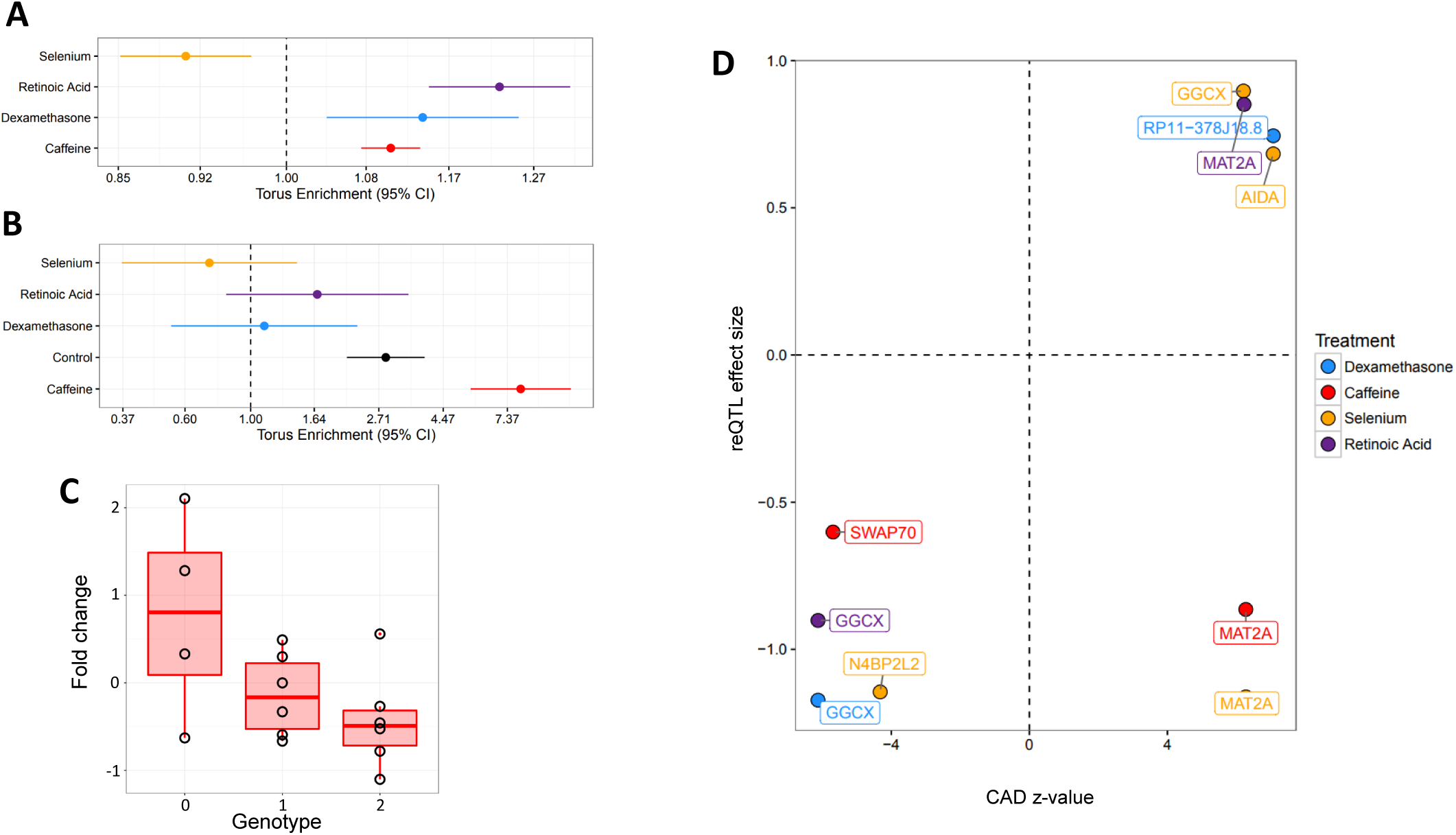
Latent environments in GTEx. **A** Enrichment of GTEx eQTLs in SNPs within response factor footprints for each treatment. **B** Enrichment of coronary artery disease risk loci in SNPs within response factor footprints for each treatment. **C** reQTL for SWAP70 in caffeine.**D** Comparison of CAD zvalue with the direction adjusted to the allele that increases gene expression vs. reQTL interaction effect with the direction adjusted to the allele that increases disease risk. A positive CAD z-value indicates that the risk allele is associated with increased gene expression in GTEx, while a negative CAD z-value indicates the risk allele is associated with decreased gene expression in GTEx. A positive reQTL effect size indicates the treatment amplifies the genetic effect on disease risk, while a negative reQTL effect size indicates the treatment reduces the genetic effect on disease risk.

### Latent environments in Coronary Artery Disease

To identify which conditions may be enriched for putative latent GxE in coronary artery disease (CAD) we performed an enrichment analysis for SNPs in footprints for response factors using CAD GWAS data [37] (Figure 3B). Among the treatment conditions we explored, we found a strikingly high enrichment of 8.17-fold for CAD associated variants that are found in caffeine response factor footprints. This enrichment is also significantly higher than the one found for footprints that are identified in the control condition (2.86-fold increase), which represent regulatory variants that do not interact with the treatments considered here. This suggests a large number of latent GxE effects in cardiovascular disease risk.

To further dissect the molecular mechanism for GxE in CAD risk, we integrated GTEx artery eQTLs with the GWAS summary statistics and GxE effects in gene expression in our data. We first performed colocalization of GTEx eQTLs with CAD GWAS hits using enloc [54] (Methods) and identified 168 SNP-gene pairs. Forty GTEx tissues had at least 1 colocalized eQTL. The most represented tissues for colocalized SNPs were the aorta and tibial artery with 56 and 28 colocalized SNPs, respectively (Supplementary Table S4). Of the colocalized genes, 14 can be tested for reQTL in our data and six have a reQTL in at least one of the treatment conditions considered (permutation p<0.05, see Supplementary TableS5). Although with our sample size we do not have the power to detect genome-wide significant signals, we provide an example of one of the top reQTL in *SWAP70* for caffeine (Figure 3C). *SWAP70* is implicated in endothelial sprouting *in vivo* [13] and the gene is significantly downregulated (BH FDR < 10%) in response to caffeine in HUVECs. The reQTL, rs10840293, is in a footprint for the caffeine response factor AFT2. These reQTLs suggest specific mechanisms through which environmental exposures can increase or decrease the genetic risk for forcoronary artery disease.

We further investigated whether the environmental interaction with the risk genotype results in amplification or buffering of the disease risk. Figure 3D shows the risk z-score from the CAD GWAS study, with a positive sign if the risk allele is associated with higher gene expression in GTEx (horizontal axis) plotted against the interaction effect in endothelial cells from our reQTL mapping. For four colocalized reQTLs, the treatment amplifies the genetic effect (positive value on the y-axis) on gene expression and in all these cases, higher expression is associated with increased risk (positive value on the x axis). For six colocalized signals, the treatment reduces the genetic effect on disease risk (negative values on the y-axis) and in most cases (4 out of 6) lower expression of the gene is associated with increased risk. For example, lower expression of *SWAP70* is associated with increased risk, however, in the presence of caffeine, the effect of the risk variant is significantly reduced (negative sign for the reQTL), thus caffeine buffers the genetic effect on disease risk. These results show that environmental effects on disease risk are contingent on the individual’s genotype, with certain exposures amplifying a specific genetic risk but buffering others.

## Discussion

Vascular endothelial dysfunction has been implicated in a variety of cardiovascular and non-cardiovascular diseases. Here we have characterized the transcriptional and chromatin landscape in endothelial cells exposed to dexamethasone, retinoic acid, caffeine, and selenium.

While all of these treatments cause significant changes in gene expression and chromatin accessibility,they act through varied pathways. Dexamethasone and retinoic acid are steroid hormones whose action at the molecular and transcriptional level has been well-studied[11, 38]. Caffeine and selenium have been investigated at the molecular level, particularly with regard to caffeine’s role in inhibiting the breakdown of cAMP and selenium’s role as a cofactor[14, 44]. However, the regulators of gene expression response to these compounds are less characterized, especially in endothelial cells. We were able to confirm with retinoic acid and dexamethasone that our approach in analyzing ATAC-seq data identifies biologically relevant response factors, as we found enrichment of known intracellular steroid receptors like the retinoic acid receptor and glucocorticoid receptor. Upon extension of this framework to caffeine and selenium, we identified key response factors for these treatments that were previously unknown (e.g. Sp1 and Sp3 for caffeine).

Combining the ATAC-seq data with known genetic variation in human populations, we have generated a set of annotations for genetic variants in the footprints of treatment response factors (1300 motifs genome-wide) which may affect binding and therefore gene expression response in the different conditions considered. Using this annotation in combination with large scale genetic studies of gene expression (GTEx) and CAD risk (Cardiogram), we detected potential latent environmental effects and assigned a putative mechanism of action for variants associated with these molecular and organismal traits, through disruption of transcription factor binding. We found that dexamethasone, retinoic acid, and caffeine were enriched in GTEx eQTLs from three different arteries in our response factor annotations. This is not surprising given the widespread use of caffeine in the general population and the expression of the glucocorticoid receptor and retinoic acid receptor in most GTEx tissues [1]. Interestingly, we did not find an enrichment for the selenium annotation, which is expected considering that we are only exposed to limited amounts of selenium on a daily basis. Our observation that that genetic loci associated with risk of coronary artery disease are highly enriched in the binding sites of transcription factors that regulate response to caffeine, suggests that caffeine modulates coronary artery disease risk. This is an intriguing finding given the extensive number of studies that have explored the link between caffeine, coffee consumption, and heart disease [12, 51].

Here, we developed a novel conceptual framework that combines information on specific environmental perturbations collected *in vitro* with large scale heterogeneous datasets interrogating genetic effects on both molecular and organismal phenotypes. While our study demonstrates that there is a significant enrichment for genetic associations in context specific regulatory annotations, our small sample size had limited power to directly validate these latent GxE effects. Nevertheless, this approach can be used to guide the design of future studies to directly test for GxE. In our proof of principle, we show that we can leverage molecular phenotyping data in response to tightly controlled treatments to characterize the complex interplay between genotypes and environment and their role in complex traits, highlighting the importance of latent environmental exposures in large-scale datasets.

## Methods

### Cell Culture

14 primary HUVECs were isolated from human umbilical cord tissue collected shortly following birth. Umbilical cord tissue specimens were obtained from healthy full-term pregnant women, admitted to DMC Hutzel Women’s Hospital (Detroit, Michigan). All specimens for this study were collected following guidelines approved by the institutional review board (#013213MP4E) of Wayne State University. Cord specimens, between 10 and 30 cm in length, were first rinsed with warm PBS and a blunt-ended needle was inserted into the umbilical vein at one end of the cord, and subsequently clamped in place. The cord was then purged to remove any excess blood from the vein. The other end of the cord was then sealed, in a manner identical to the first end, and pre-warmed 0.25% trypsin-EDTA (Gibco) was then injected into the vein. Following a 20 minute incubation, at 37°C, detached HUVECs were rinsed from the vein, collected by centrifugation, counted, and seeded into an appropriate vessel at 10,000 cells/cm^2^, in EGM-2 growth medium (Lonza). Expanded cultures were cryopreserved prior to be used in the experiments. For additional 3 HUVEC lines from a previous study [35] we collected ATAC-seq data and used previously published RNA-seq data (dbGaP accession number phs001176.v1.p1).

### Treatments

Treatment concentrations were the same as used in [35] and were derived from the Clinical Guidelines Mayo Clinic Reference Levels (http://www.mayomedicallaboratories.com) and the CDC National Biomonitoring Report Reference Levels (http://www.cdc.gov/biomonitoring/). Concentrations were 1 10^−6^ M for dexamethasone, 1×10^−8^ M for retinoic acid, 1.16×10^−3^ M for caffeine, and 1×10^−5^ M for selenium. Vehicle controls were included to represent the solvent used to prepare the different treatments, either ethanol (1*μ*l of EtOH per 10,000*μ*l of culturing media) or water.

Each individual cell line was treated on a separate day. Culturing conditions were the same as those found in [35]. Specifically, three days before treatment, cells were plated (to passage 7) in 8 wells in 6-well plates at 5,000/cm^2^ in EGM-2. Following a 24 hour recovery period, the medium was changed to a “starvation medium” composed of phenol-red free EGM-2, without Hydrocortisone and Vitamin C and supplemented with 2% CS-FBS. Cell starvation was continued for 48 hour prior to treatment. For each individual cell line, control treatments were performed in duplicate. Cells were treated for 6 hours and then were collected by scraping the plate on ice. For four cell lines, each well was counted in order to remove 50,000 cells to be used for ATAC-seq while the rest were used for RNA sequencing.

### RNA Library Preparation

Treated cells were collected by centrifugation at 2000 rpm and washed 2x using ice cold PBS. Collected pellets were lysed on the plate, using Lysis/Binding Buffer (Ambion), and frozen at −80°. Poly-adenylated mRNAs were subsequently isolated from thawed lysates using the Dynabeads mRNA Direct Kit (Ambion) and following the manufacturer instructions. RNA-seq libraries were prepared using a protocol modified from the NEBNext Ultradirectional (NEB) library preparation protocol to use 96 Barcodes from BIOO-Scientific added by ligation, as described in [33]. The individual libraries were quantified using the KAPA real-time PCR system, following the manufacturer instructions and using a custom-made series of standards obtained from serial dilutions of the phi-X DNA (Illumina). Libraries were pooled and sequenced in multiple sequencing runs for an average of 50M 300bp PE reads.

### ATAC-seq Library Preparation

We followed the protocol by [7] to lyse 50,000 cells and prepare ATAC-seq libraries, with the exception that we used the Illumina Nextera Index Kit (Cat #15055290) in the PCR enrichment step. For 3 of the 4 individual cell lines (LP-001, LP-002 and H288-L), the cells were not lysed with 0.1% IGEPAL CA-630 before adding the transposase to begin the ATAC-seq protocol. Individual library fragment distributions were assessed on the Agilent Bioanalyzer and pooling proportions were determined using the qPCR Kapa library quantification kit (KAPA Biosystems). Library pools were run on the Illumina NextSeq 500 Desktop sequencer in the Luca/Pique-Regi laboratory. Barcoded libraries of ATAC-seq samples were pooled and sequenced in multiple sequencing runs for an average coverage of 130M reads.

### Alignment of RNA and ATAC-seq

Reads were aligned to the GRCh37 human reference genome using HISAT2 [26] (https://ccb.jhu.edu/software/hisat2/index.shtml, version hisat2-2.0.4), and the human reference genome(GRCh37) with the following options:

~~~
HISAT2 -x <genome> −1 <fastq_R1.gz> −2 <fastq_R2.gz>
~~~

where `<genome>` represents the location of the genome file (genome snp for ATAC-seq alignment, genome snp tran for RNA-seq alignment), and `<fastqs R1.gz>` and `<fastqs R2.gz>` represent that sample’s fastq files.

The multiple sequencing runs were merged for each samples using samtools (version 2.25.0). We removed PCR duplicates and further removed reads with a quality score of < 10 (equating to reads mapped to multiple locations) for the RNA-sequencing analysis, only.

### Differential Gene Expression Analysis

To identify differentially expressed (DE) genes, we used `DESeq2` [30] (R version 3.2.1, `DESeq2` version 1.8.1). Gene annotations from Ensembl version 75 were used and transcripts with fewer than 20 reads total were discarded. coverageBed was utilized to count reads with -s to account for strandedness and -split for BED12 input. The counts were then utilized in DESeq2 to determine changes in gene expression under the different treatment conditions. Multiple test correction was performed using the Benjamini-Hochberg procedure [3] with a significance threshold of 10%. A gene was considered DE if at least one of its transcripts was DE and had an absolute log_2_ fold change *>* 0.25. The model corrected for the library preparation batch and the sample of origin.

### Gene ontology analysis

For each treatment, we tested upregulated and downregulated genes separately for enrichment of biological processes gene ontology terms using the Fisher’s test on the AmiGO 2 web browser (http://amigo.geneontology.org/amigo). Multiple test correction was performed using the Benjamini-Hochberg procedure with a significance threshold of 5% (BH-FDR).

### Identification of Differentially Accessible Regions Following Treatment

To identify differentially accessible regions, we used `DESeq2` [30] (R version 3.2.1, `DESeq2` version 1.8.1). We separated the genome into 300bp regions and `coverageBed` was used to count reads in these regions. A total of 508,140 regions with *>* 0.25 reads per million (high accessibility regions) were then utilized in DESeq2 using the following model:

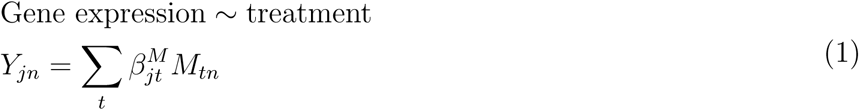

where *Y_jn_* represents the internal DESeq2 mean accessibility parameter for region *j* and experiment *n*,*M_n_* is the treatment indicator, and 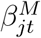 parameter is the treatment effect. Differentially accessible regions (DAR) following exposure to dexamethasone, retinoic acid, caffeine, and selenium, were determined at BH FDR < 10%. We annotated the genomic context of these DARs using chromHMM [16], which characterizes chromatin states on the basis of histone marks. We downloaded the precomputed HUVEC annotations from ENCODE [15]. Enrichments were performed with the Fisher’s Exact test. DARs were then compared to gene annotations from Ensembl version 75 to identify those that were within 50kb of a transcription start site, and a Fisher’s exact test was used to test for enrichment of DARs near DE genes.

### Transcription factor binding footprints

To detect which transcription factors have footprints in each condition we adapted CENTIPEDE [41] to use the fragment length information contained in the ATAC-seq in the footprint model, and to jointly use in parallel the treatment and control conditions in order to ensure that the same footprint shape is used for the same motif in both conditions.

As in CENTIPEDE, we need to start from candidate binding sites for a given motif model. For each transcription factor we scan the entire human genome (hg19) for matches to its DNA recognition motif using position weight matrix (PWM) models from TRANSFAC and JASPAR as previously described[41]. Then for each candidate location *l* we collect all the ATAC-seq fragments which are partitioned into four binds depending on the fragment length: 1) [39-99], 2) [100-139], 3) [140-179], 4) [180-250]. For each fragment, the two Tn5 insertion sites were calculated as the position 4bp after the 5’-end in the 5’ to 3’ direction. Then for each candidate motif, a matrix ***X*** was constructed to count Tn5 insertion events: each row represented a sequence match to motif in the genome (motif instance), and each column a specific cleavage site at a relative bp and orientation with respect to the motif instance. We built a matrix 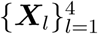 for each fragment length bin, each using a window half-size S=150bp resulting in (2 *S* + *W*)×2 columns, where W is the length of the motif in bp.

Finally, we fit the CENTIPEDE model in a subset of approximately 5,000 instances to learn footprint shapes for each factor as in [34]. Then we used CENTIPEDE to analyze the remaining motif instances in the genome and kept those with a CENTIPEDE posterior probability higher than 0.99 to denote locations where the transcription factors are bound also referred as ‘footprints’. We further subset these bound transcription factor sites in each condition by including only transcription factors which were active genome-wide, as indicated by an enrichment of binding sites in high accessibility regions (see above) compared to a randomly selected set of 500,000 chromatin regions genome-wide (not high accessible regions).

### Identification of response factors and genetic variants in their binding sites

In order to identify transcription factors which are important for response to treatments, we calculated enrichment scores for each active motif in regions of differentially accessible chromatin using a Fisher’s exact test. We defined motifs that were significantly enriched or depleted in opening or closing chromatin (nominal *p*-value < 0.05, Table S3). For each tissue and in each treatment, we then annotated each SNP from the 1000 Genomes project[20] as 1) not in a footprint for a treatment response factor; 2) in a footprint for a treatment response factor (Supplementary Table S2). SNPs in footprints are generally enriched in eQTLs [53]. Thus, to account for this treatment-independent effect, we also generated an annotation for SNPs in active footprints from the control condition, which should represent a baseline annotation of candidate regulatory variants.

### Enrichment of GTEx eQTL and CAD variants in footprints for response factors

To identify treatments for which GTEx eQTLs[1] are enriched in footprints for response factors, we focused on the three artery tissues present in GTEx: tibial, coronary, and aorta. GTEx identified 11,945, 4,378, and 9,203 genes with an eQTL (eGenes) in the tibial artery, coronary artery, and aorta tissues, respectively. We then used the Torus package [53] with the `-est` option to calculate the enrichment of eQTLs in response factors for each treatment. Torus quantitatively assesses the enrichment of molecular annotations (e.g. SNPs in response factors) in GWAS or eQTL datasets. Here we used the annotations generated above for SNPs in footprints for response factors and control. As additional annotation, we included distance of the SNP to the transcriptional start site of the gene. We meta-analyzed the enrichments for each treatment across tissues using a fixed effects model with inverse variance weights. For the CAD enrichment, we used the same annotations as above on the summary statistics of the CAD GWAS, minus the distance of the SNP to the transcriptional start site of a gene. GWAS SNPs were partitioned into blocks based on LD pattern.

### Colocalization

We performed colocalization as previously described using *enloc*[54]. Briefly, *enloc* integrates QTL annotations into GWAS analysis by calculating the enrichment of QTLs in complex trait-associated genetic variants, which is then used to identify colocalized SNPs between the two annotations. We performed the colocalization between the CARDIoGRAM CAD summary statistics and eQTLs from all GTEx tissues independently. We considered the lead SNP from a locus with a cumulative posterior inclusion probability (PIP) of greater than 0.5 to be colocalized. 168 SNPs colocalize, corresponding to 139 genes.

DNA was isolated from umbilical cords and sent to be genotyped using ultra-low sequencing performed by Gencove. In our dataset, 21 of the colocalized SNPs (corresponding to 14 eGenes) are polymorphic and therefore can be tested for reQTL.

### reQTL mapping

For reQTL mapping of the colocalized GTEx and CAD GWAS SNPs, we tested all 21 SNPs for reQTLs, corresponding to 33 gene-SNP pairs. We used a linear model to identify reQTLs, testing for the effect of genotype dosage on quantile-normalized gene expression fold change between treatment and control. We generated an empirical null distribution by permuting sample labels, and corrected the treatment p-values based on the ranks observed in the permutation. We performed 2,820 permutations, corresponding to 3x the number of tests in the treated samples.

### Accession numbers for sequencing data

Data upload to dbGaP ID phs001176.v1.p1 is pending.

### Additional Files

See supplementary tables and figures for additional results.

## Acknowledgements

We thank members of the Luca/Pique-Regi group for helpful discussions and comments. This work was supported by the American Heart Association (14SDG20450118 to F.L.), the National Institute of General Medical Sciences of the National Institutes of Health (R01GM109215 to F.L. and R.P.).

## Supplementary Materials

### Supplementary Tables

**Table S1: Gene ontology results.** Gene ontology output for up- and down-regulated genes. Note that retinoic acid and selenium had no significantly enriched gene ontology terms for upregulated genes.

A) http://genome.grid.wayne.edu/FUNGEI/data_tables/Caffeine_Down.txt

B) http://genome.grid.wayne.edu/FUNGEI/data_tables/Caffeine_Up.txt

C) http://genome.grid.wayne.edu/FUNGEI/data_tables/Dexamethasone_Down.txt

D) http://genome.grid.wayne.edu/FUNGEI/data_tables/Dexamethasone_Up.txt E)http://genome.grid.wayne.edu/FUNGEI/data_tables/Retinoic_Acid_Down.txt

F) http://genome.grid.wayne.edu/FUNGEI/data_tables/Selenium_Down.txt

**Table S2: 1000 Genomes SNP annotations.** We have annotated 1000 Genomes SNPs on the basis of whether they are within (1) or not within (0) a response factor footprint for each treatment. Column 1 is the SNP ID including chromosome and position, column 2 - 6 are annotations for dexamethasone, caffeine, selenium, retinoic acid, and control, respectively.

http://genome.grid.wayne.edu/FUNGEI/data_tables/FUNGEI_anno.v2.combined_1_2.active.gz

**Table S3: Enrichment of footprints in DARs.** For retinoic acid (A), dexamethasone (B), caffeine (C), and selenium (D), we have included the motif enrichment in DARs in both opening and closing chromatin. Columns 1-18 are: 1) Motif ID, 2) Transcription factor name, 3) Odds ratio from Fisher’s exact test for opening DARs, 4) Lower bound of 95% confidence interval of odds ratio for opening DARs, 5) Upper bound of 95% confidence interval of odds ratio for opening DARs, 6) P-value from Fisher’s exact test for opening DARs, 7) Number of footprints in opening DARs, 8) Number of footprints in regions tested for differential accessibility that are not opening DARs, 9) Number of opening DARs without footprint, 10) Number of regions tested for differential accessibility without opening DARs and without footprint, 11) Odds ratio from Fisher’s exact test for closing DARs, 12) Lower bound of 95% confidence interval of odds ratio for closing DARs, 13) Upper bound of 95% confidence interval of odds ratio for closing DARs, 14) P-value from Fisher’s exact test for closing DARs, 15) Number of footprints in closing DARs, 16) Number of footprints in regions tested for differential accessibility that are not closing DARs, Number of closing DARs without footprint, 18) Number of regions tested for differential accessibility without closing DARs and without footprint

A) http://genome.grid.wayne.edu/FUNGEI/data_tables/Retinoic_Acid_all_Fisher.tab

B) http://genome.grid.wayne.edu/FUNGEI/data_tables/Dexamethasone_all_Fisher.tab

C) http://genome.grid.wayne.edu/FUNGEI/data_tables/Caffeine_all_Fisher.tab

D) http://genome.grid.wayne.edu/FUNGEI/data_tables/Selenium_all_Fisher.tab

**Table S4: Colocalization results.** This file includes the output from the colocalization with enloc. Columns 1-9 are: 1) eQTL signal ID 2) eQTL posterior inclusion probability (PIP) 3) GWAS PIP 4) number of SNPs in the group 5) original enloc regional colocalization probability (RCP) 6) experimental enloc PIP 7) lead SNP in the signal group 8) SNP-level colocalization probability for the lead SNP 9) tissue.

http://genome.grid.wayne.edu/FUNGEI/data_tables/Torus_output_tissues.txt

**Table S5: Colocalization reQTLs.** This file includes reQTLs with permutation p<0.05. Columns 1-6 are: 1) rsID of reQTL, 2) gene ID, 3) treatment (sel = selenium, dex = dexamethasone, ret = retinoic acid, caf = caffeine),4) reQTL effect size, 5) permutation p-value, 6) zvalue from coronary artery disease (CAD) GWAS.

http://genome.grid.wayne.edu/FUNGEI/data_tables/CAD_reQTLs.txt

**Table S6: Active factor enrichment.** This file includes the enrichment of footprints of each motif in each condition in highly active chromatin. Columns 1-17 are: 1) Motif ID, 2) Transcription factor name, 3) Odds ratio of enrichment of footprints in dexamethasone in highly accessible regions, 4) P-value of enrichment of footprints in dexamethasone in highly accessible regions, 5) Benjamini-Hochberg adjusted p-value of enrichment of footprints in dexamethasone in highly accessible regions, 6) Odds ratio of enrichment of footprints in caffeine in highly accessible regions, 7) P-value of enrichment of footprints in caffeine in highly accessible regions, 8) Benjamini-Hochberg adjusted p-value of enrichment of footprints in caffeine in highly accessible regions, 9) Odds ratio of enrichment of footprints in selenium in highly accessible regions, 10) P-value of enrichment of footprints in selenium in highly accessible regions, 11) Benjamini-Hochberg adjusted p-value of enrichment of footprints in selenium in highly accessible regions, 12) Odds ratio of enrichment of footprints in retinoic acid in highly accessible regions, 13) P-value of enrichment of footprints in retinoic acid in highly accessible regions, 14) Benjamini-Hochberg adjusted p-value of enrichment of footprints in retinoic acid in highly accessible regions, 15) Odds ratio of enrichment of footprints in control condition in highly accessible regions, 16) P-value of enrichment of footprints in control condition in highly accessible regions, 17) Benjamini-Hochberg adjusted p-value of enrichment of footprints in control condition in highly accessible regions.

http://genome.grid.wayne.edu/FUNGEI/data_tables/all.active_factor_enrichment.txt

---

**Figure S1:**
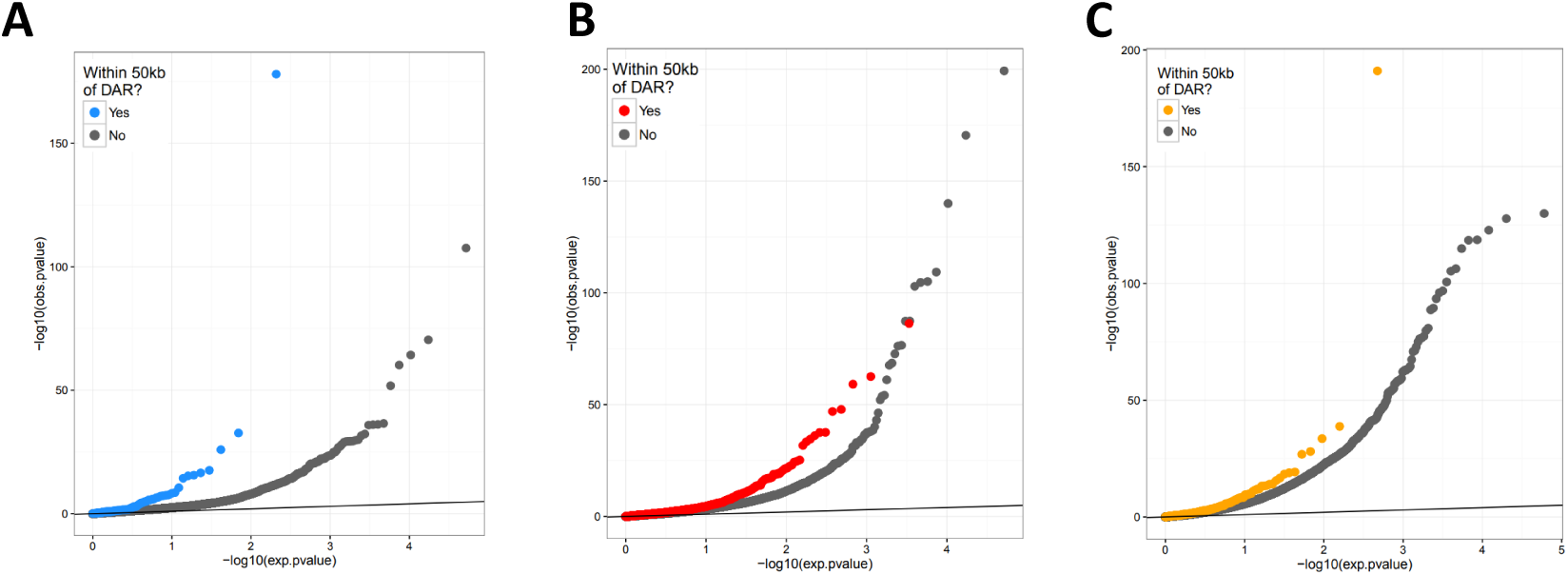
Differential gene expression near DARs. QQ-plot of *p*-values from DESeq2 analysis of gene expression for dexamethasone (**A**), caffeine (**B**), and selenium (**C**). The colored dots represent genes within 50kb of a DAR, while the gray dots represent genes not within 50kb of a DAR..

**Figure S2:**
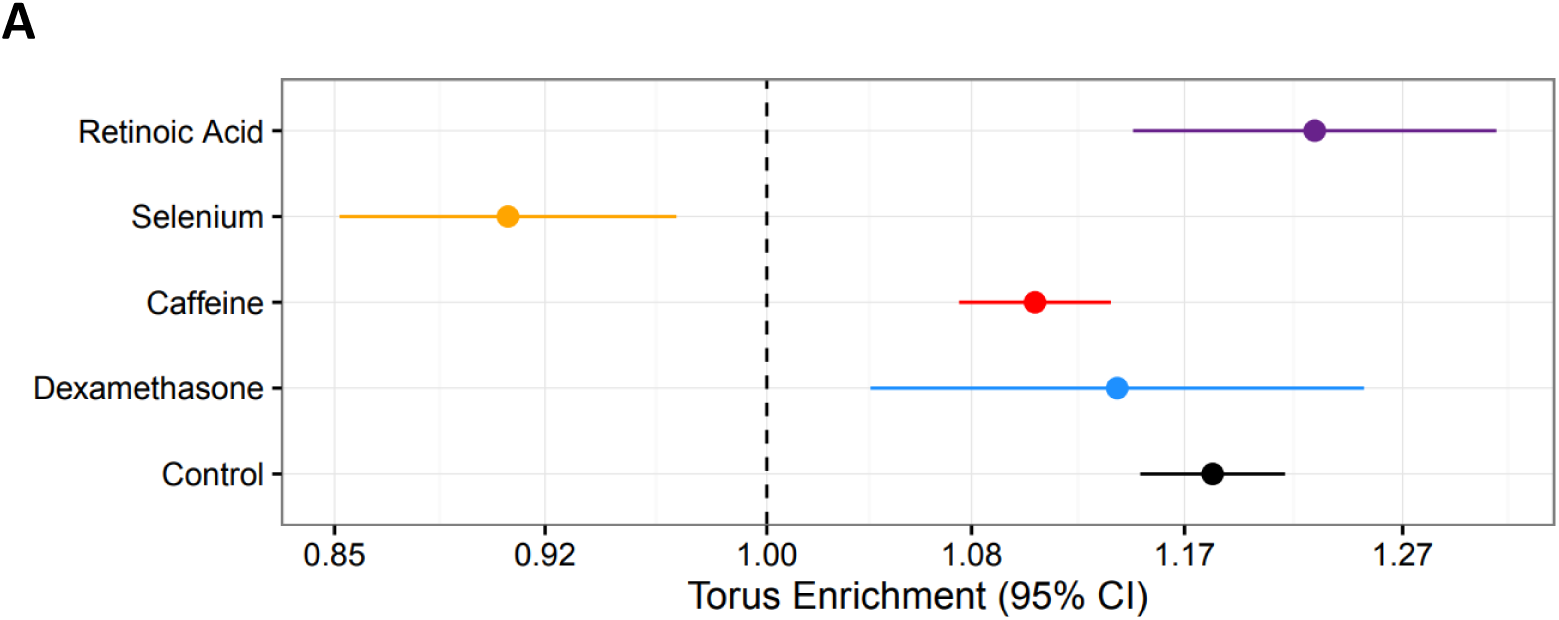
Torus enrichment of response factor SNPs in GTEx Tibial Artery eQTLs. Enrichment of GTEx eQTLs in SNPs within response factor footprints in each treatment. This is identical to Figure 3A, but includes the enrichment in the control condition.

## References

[1] François Aguet, Andrew A. Brown, Stephane E. Castel, Joe R. Davis, Yuan He, Brian Jo, Pejman Mohammadi, YoSon Park, Princy Parsana, Ayellet V. Segré, Benjamin J. Strober, Zachary Zappala, Beryl B. Cummings, Ellen T. Gelfand, Kane Hadley, Katherine H. Huang, Monkol Lek, Xiao Li, Jared L. Nedzel, Duyen Y. Nguyen, Michael S. Noble, Timothy J. Sullivan, Taru Tukiainen, Daniel G. MacArthur, Gad Getz, Anjene Addington, Ping Guan, Susan Koester, A. Roger Little, Nicole C. Lockhart, Helen M. Moore, Abhi Rao, Jeffery P. Struewing, Simona Volpi, Lori E. Brigham, Richard Hasz, Marcus Hunter, Christopher Johns, Mark Johnson, Gene Kopen, William F. Leinweber, John T. Lonsdale, Alisa McDonald, Bernadette Mestichelli, Kevin Myer, Bryan Roe, Michael Salvatore, Saboor Shad, Jeffrey A. Thomas, Gary Walters, Michael Washington, Joseph Wheeler, Jason Bridge, Barbara A. Foster, Bryan M. Gillard, Ellen Karasik, Rachna Kumar, Mark Miklos, Michael T. Moser, Scott D. Jewell, Robert G. Montroy, Daniel C. Rohrer, Dana Valley, Deborah C. Mash, David A. Davis, Leslie Sobin, Mary E. Barcus, Philip A. Branton, Nathan S. Abell, Brunilda Balliu, Olivier Delaneau, Laure Frésard, Eric R. Gamazon, Diego Garrido-Martín, Ariel D. H. Gewirtz, Genna Gliner, Michael J. Gloudemans, Buhm Han, Amy Z. He, Farhad Hormozdiari, Xin Li, Boxiang Liu, Eun Yong Kang, Ian C. McDowell, Halit Ongen, John J. Palowitch, Christine B. Peterson, Gerald Quon, Stephan Ripke, Ashis Saha, Andrey A. Shabalin, Tyler C. Shimko, Jae Hoon Sul, Nicole A. Teran, Emily K. Tsang, Hailei Zhang, Yi-Hui Zhou, Carlos D. Bustamante, Nancy J. Cox, Roderic Guigó, Manolis Kellis, Mark I. McCarthy, Donald F. Conrad, Eleazar Eskin, Gen Li, Andrew B. Nobel, Chiara Sabatti, Barbara E. Stranger, Xiaoquan Wen, Fred A. Wright, Kristin G. Ardlie, Emmanouil T. Dermitzakis, Tuuli Lappalainen, François Aguet, Kristin G. Ardlie, Beryl B. Cummings, Ellen T. Gelfand, Gad Getz, Kane Hadley, Robert E. Handsaker, Katherine H. Huang, Seva Kashin, Konrad J. Karczewski, Monkol Lek, Xiao Li, Daniel G. MacArthur, Jared L. Nedzel, Duyen T. Nguyen, Michael S. Noble, Ayellet V. Segrè, Casandra A. Trowbridge, Taru Tukiainen, Nathan S. Abell, Brunilda Balliu, Ruth Barshir, Omer Basha, Alexis Battle, Gireesh K. Bogu, Andrew Brown, Christopher D. Brown, Stephane E. Castel, Lin S. Chen, Colby Chiang, Donald F. Conrad, Nancy J. Cox, Farhan N. Damani, Joe R. Davis, Olivier Delaneau, Emmanouil T. Dermitzakis, Barbara E. Engelhardt, Eleazar Eskin, Pedro G. Ferreira, Laure Frésard, Eric R. Gamazon, Diego Garrido-Martín, Ariel D. H. Gewirtz, Genna Gliner, Michael J. Gloudemans, Roderic Guigo, Ira M. Hall, Buhm Han, Yuan He, Farhad Hormozdiari, Cedric Howald, Hae Kyung Im, Brian Jo, Eun Yong Kang, Yungil Kim, Sarah Kim-Hellmuth, Tuuli Lappalainen, Gen Li, Xin Li, Boxiang Liu, Serghei Mangul, Mark I. McCarthy, Ian C. McDowell, Pejman Mohammadi, Jean Monlong, Stephen B. Montgomery, Manuel Muñoz-Aguirre, Anne W. Ndungu, Dan L. Nicolae, Andrew B. Nobel, Meritxell Oliva, Halit Ongen, John J. Palowitch, Nikolaos Panousis, Panagiotis Papasaikas, YoSon Park, Princy Parsana, Anthony J. Payne, Christine B. Peterson, Jie Quan, Ferran Reverter, Chiara Sabatti, Ashis Saha, Michael Sammeth, Alexandra J. Scott, Andrey A. Shabalin, Reza Sodaei, Matthew Stephens, Barbara E. Stranger, Benjamin J. Strober, Jae Hoon Sul, Emily K. Tsang, Sarah Urbut, Martijn van de Bunt, Gao Wang, Xiaoquan Wen, Fred A. Wright, Hualin S. Xi, Esti Yeger-Lotem, Zachary Zappala, Judith B. Zaugg, Yi-Hui Zhou, Joshua M. Akey, Daniel Bates, Joanne Chan, Lin S. Chen, Melina Claussnitzer, Kathryn Demanelis, Morgan Diegel, Jennifer A. Doherty, Andrew P. Feinberg, Marian S. Fernando, Jessica Halow, Kasper D. Hansen, Eric Haugen, Peter F. Hickey, Lei Hou, Farzana Jasmine, Ruiqi Jian, Lihua Jiang, Audra Johnson, Rajinder Kaul, Manolis Kellis, Muhammad G. Kibriya, Kristen Lee, Jin Billy Li, Qin Li, Xiao Li, Jessica Lin, Shin Lin, Sandra Linder, Caroline Linke, Yaping Liu, Matthew T. Maurano, Benoit Molinie, Stephen B. Montgomery, Jemma Nelson, Fidencio J. Neri, Meritxell Oliva, Yongjin Park, Brandon L. Pierce, Nicola J. Rinaldi, Lindsay F. Rizzardi, Richard Sandstrom, Andrew Skol, Kevin S. Smith, Michael P. Snyder, John Stamatoyannopoulos, Barbara E. Stranger, Hua Tang, Emily K. Tsang, Li Wang, Meng Wang, Nicholas Van Wittenberghe, Fan Wu, Rui Zhang, Concepcion R. Nierras, Philip A. Branton, Latarsha J. Carithers, Ping Guan, Helen M. Moore, Abhi Rao, Jimmie B. Vaught, Sarah E. Gould, Nicole C. Lockart, Casey Martin, Jeffery P. Struewing, Simona Volpi, Anjene M. Addington, Susan E. Koester, A. Roger Little, Lori E. Brigham, Richard Hasz, Marcus Hunter, Christopher Johns, Mark Johnson, Gene Kopen, William F. Leinweber, John T. Lonsdale, Alisa McDonald, Bernadette Mestichelli, Kevin Myer, Brian Roe, Michael Salvatore, Saboor Shad, Jeffrey A. Thomas, Gary Walters, Michael Washington, Joseph Wheeler, Jason Bridge. Genetic effects on gene expression across human tissues. Nature, 550(7675):204–213, oct 2017.

[2] Kaur Alasoo, Julia Rodrigues, Subhankar Mukhopadhyay, Andrew J Knights, Alice L Mann, Kousik Kundu, Christine HIPSCI Consortium, Christine Hale, Gordon Dougan, and Daniel J Gaffney. Shared genetic effects on chromatin and gene expression indicate a role for enhancer priming in immune response. Nature genetics, 50(3):424–431, mar 2018.

[3] Yoav Benjamini and Yosef Hochberg. Controlling the False Discovery Rate: A Practical and Powerful Approach to Multiple Testing. Journal of the Royal Statistical Society. Series B (Methodological), 57(1):289–300, 1995.

[4] Ana R. Bernardo, José M. Cosgaya, Ana Aranda, and Ana M. Jiménez-Lara. Pro-apoptotic signaling induced by Retinoic acid and dsRNA is under the control of Interferon Regulatory Factor-3 in breast cancer cells. Apoptosis, 22(7):920–932, jul 2017.

[5] Alan P Boyle, Sean Davis, Hennady P Shulha, Paul Meltzer, Elliott H Margulies, Zhiping Weng, Terrence S Furey, and Gregory E Crawford. High-resolution mapping and characterization of open chromatin across the genome. Cell, 132(2):311–22, jan 2008.

[6] Jason D Buenrostro, Paul G Giresi, Lisa C Zaba, Howard Y Chang, and William J Greenleaf. Transposition of native chromatin for fast and sensitive epigenomic profiling of open chromatin, DNA-binding proteins and nucleosome position. Nat. Methods, (October):1–8, oct 2013.

[7] Jason D Buenrostro, Paul G Giresi, Lisa C Zaba, Howard Y Chang, and William J Greenleaf. Transposition of native chromatin for fast and sensitive epigenomic profiling of open chromatin, DNA-binding proteins and nucleosome position. Nature methods, 10(12):1213–8, dec 2013.

[8] Krysta Mila Coyle, Selena Maxwell, Margaret Lois Thomas, and Paola Marcato. Profiling of the transcriptional response to all-trans retinoic acid in breast cancer cells reveals RARE-independent mechanisms of gene expression. Scientific Reports, 7(1):16684, dec 2017.

[9] Jacob F Degner, Athma a Pai, Roger Pique-Regi, Jean-Baptiste B Veyrieras, Daniel J Gaffney, Joseph K Pickrell, Sherryl De Leon, Katelyn Michelini, Noah Lewellen, Gregory E Crawford, Matthew Stephens, Yoav Gilad, and Jonathan K Pritchard. DNase I sensitivity QTLs are a major determinant of human expression variation. Nature, 482(7385):390–394, feb 2012.

[10] E Deniaud, J Baguet, A-L Mathieu, G Pagès, J Marvel, and Y Leverrier. Overexpression of Sp1 transcription factor induces apoptosis. Oncogene, 25(53):7096–105, nov 2006.

[11] Alessandra di Masi, Loris Leboffe, Elisabetta De Marinis, Francesca Pagano, Laura Cicconi, Cécile Rochette-Egly, Francesco Lo-Coco, Paolo Ascenzi, and Clara Nervi. Retinoic acid receptors: From molecular mechanisms to cancer therapy. Molecular Aspects of Medicine, 41:1–115, feb 2015.

[12] Ming Ding, Shilpa N Bhupathiraju, Ambika Satija, Rob M van Dam, and Frank B Hu. Long-term coffee consumption and risk of cardiovascular disease: a systematic review and a dose-response metaanalysis of prospective cohort studies. Circulation, 129(6):643–59, feb 2014.

[13] Julie Dwyer, Sandy Azzi, Héloïse M Leclair, Steven Georges, Agnès Carlotti, Lucas Treps, Eva M Galan-Moya, Catherine Alexia, Nicolas Dupin, Nicolas Bidère, and Julie Gavard. The guanine exchange factor SWAP70 mediates vGPCR-induced endothelial plasticity. Cell Communication and Signaling, 13(1):11, feb 2015.

[14] Darío Echeverri, Félix R Montes, Mariana Cabrera, Angélica Galán, and Angélica Prieto. Caffeine’s Vascular Mechanisms of Action. International journal of vascular medicine, 2010:834060, 2010.

[15] ENCODE Project Consortium. An integrated encyclopedia of DNA elements in the human genome. Nature, 489(7414):57–74, sep 2012.

[16] Jason Ernst and Manolis Kellis. ChromHMM: automating chromatin-state discovery and characterization. Nature Methods, 9(3):215–216, mar 2012.

[17] Gemma Flores-Mateo, Ana Navas-Acien, Roberto Pastor-Barriuso, and Eliseo Guallar. Selenium and coronary heart disease: a meta-analysis. The American Journal of Clinical Nutrition, 84(4):762–773, oct 2006.

[18] Eric R Gamazon, Heather E Wheeler, Kaanan P Shah, Sahar V Mozaffari, Keston Aquino-Michaels, Robert J Carroll, Anne E Eyler, Joshua C Denny, Dan L Nicolae, Nancy J Cox, Hae Kyung Im, and Hae Kyung Im. A gene-based association method for mapping traits using reference transcriptome data. Nature Genetics, 47(9):1091–1098, sep 2015.

[19] Rachel E Gate, Christine S Cheng, Aviva P Aiden, Atsede Siba, Marcin Tabaka, Dmytro Lituiev, Ido Machol, M Grace Gordon, Meena Subramaniam, Muhammad Shamim, Kendrick L Hougen, Ivo Wortman, Su-Chen Huang, Neva C Durand, Ting Feng, Philip L De Jager, Howard Y Chang, Erez Lieberman Aiden, Christophe Benoist, Michael A Beer, Chun J Ye, and Aviv Regev. Genetic determinants of co-accessible chromatin regions in activated T cells across humans. Nature genetics, 50(8):1140–1150, aug 2018.

[20] Richard A. Gibbs, Eric Boerwinkle, Harsha Doddapaneni, Yi Han, Viktoriya Korchina, Christie Kovar, Sandra Lee, Donna Muzny, Jeffrey G. Reid, Yiming Zhu, Jun Wang, Yuqi Chang, Qiang Feng, Xiaodong Fang, Xiaosen Guo, Min Jian, Hui Jiang, Xin Jin, Tianming Lan, Guoqing Li, Jingxiang Li, Yingrui Li, Shengmao Liu, Xiao Liu, Yao Lu, Xuedi Ma, Meifang Tang, Bo Wang, Guangbiao Wang, Honglong Wu, Renhua Wu, Xun Xu, Ye Yin, Dandan Zhang, Wenwei Zhang, Jiao Zhao, Meiru Zhao, Xiaole Zheng, Eric S. Lander, David M. Altshuler, Stacey B. Gabriel, Namrata Gupta, Neda Gharani, Lorraine H. Toji, Norman P. Gerry, Alissa M. Resch, Paul Flicek, Jonathan Barker, Laura Clarke, Laurent Gil, Sarah E. Hunt, Gavin Kelman, Eugene Kulesha, Rasko Leinonen, William M. McLaren, Rajesh Radhakrishnan, Asier Roa, Dmitriy Smirnov, Richard E. Smith, Ian Streeter, Anja Thormann, Iliana Toneva, Brendan Vaughan, Xiangqun Zheng-Bradley, David R. Bentley, Russell Grocock, Sean Humphray, Terena James, Zoya Kingsbury, Hans Lehrach, Ralf Sudbrak, Marcus W. Albrecht, Vyacheslav S. Amstislavskiy, Tatiana A. Borodina, Matthias Lienhard, Florian Mertes, Marc Sultan, Bernd Timmermann, Marie-Laure Yaspo, Elaine R. Mardis, Richard K. Wilson, Lucinda Fulton, Robert Fulton, Stephen T. Sherry, Victor Ananiev, Zinaida Belaia, Dimitriy Beloslyudtsev, Nathan Bouk, Chao Chen, Deanna Church, Robert Cohen, Charles Cook, John Garner, Timothy Hefferon, Mikhail Kimelman, Chunlei Liu, John Lopez, Peter Meric, Chris O’Sullivan, Yuri Ostapchuk, Lon Phan, Sergiy Ponomarov, Valerie Schneider, Eugene Shekhtman, Karl Sirotkin, Douglas Slotta, Hua Zhang, Gil A. McVean, Richard M. Durbin, Senduran Balasubramaniam, John Burton, Petr Danecek, Thomas M. Keane, Anja Kolb-Kokocinski, Shane McCarthy, James Stalker, Michael Quail, Jeanette P. Schmidt, Christopher J. Davies, Jeremy Gollub, Teresa Webster, Brant Wong, Yiping Zhan, Adam Auton, Christopher L. Campbell, Yu Kong, Anthony Marcketta, Richard A. Gibbs, Fuli Yu, Lilian Antunes, Matthew Bainbridge, Donna Muzny, Aniko Sabo, Zhuoyi Huang, Jun Wang, Lachlan J. M. Coin, Lin Fang, Xiaosen Guo, Xin Jin, Guoqing Li, Qibin Li, Yingrui Li, Zhenyu Li, Haoxiang Lin, Binghang Liu, Ruibang Luo, Haojing Shao, Yinlong Xie, Chen Ye, Chang Yu, Fan Zhang, Hancheng Zheng, Hongmei Zhu, Can Alkan, Elif Dal, Fatma Kahveci, Gabor T. Marth, Erik P. Garrison, Deniz Kural, Wan-Ping Lee, Wen Fung Leong, Michael Stromberg, Alistair N. Ward, Jiantao Wu, Mengyao Zhang, Mark J. Daly, Mark A. DePristo, Robert E. Handsaker, David M. Altshuler, Eric Banks, Gaurav Bhatia, Guillermo del Angel, Stacey B. Gabriel, Giulio Genovese, Namrata Gupta, Heng Li, Seva Kashin, Eric S. Lander, Steven A. McCarroll, James C. Nemesh, Ryan E. Poplin, Seungtai C. Yoon, Jayon Lihm, Vladimir Makarov, Andrew G. Clark, Srikanth Gottipati, Alon Keinan, Juan L. Rodriguez-Flores, Jan O. Korbel, Tobias Rausch, Markus H. Fritz, Adrian M. Stütz, Paul Flicek, Kathryn Beal, Laura Clarke, Avik Datta, Javier Herrero, William M. McLaren, Graham R. S. Ritchie, Richard E. Smith, Daniel Zerbino, Xiangqun Zheng-Bradley, Pardis C. Sabeti, Ilya Shlyakhter, Stephen F. Schaffner, Joseph Vitti, David N. Cooper, Edward V. Ball, Peter D. Stenson, David R. Bentley, Bret Barnes, Markus Bauer, R. Keira Cheetham, Anthony Cox, Michael Eberle, Sean Humphray, Scott Kahn, Lisa Murray, John Peden, Richard Shaw, Eimear E. Kenny, Mark A. Batzer, Miriam K. Konkel, Jerilyn A. Walker, Daniel G. MacArthur, Monkol Lek, Ralf Sudbrak, Vyacheslav S. Amstislavskiy, Ralf Herwig, Elaine R. Mardis, Li Ding, Daniel C. Koboldt, David Larson, Kai Ye, Simon Gravel, Anand Swaroop, Emily Chew, Tuuli Lappalainen, Yaniv Erlich, Melissa Gymrek, Thomas Frederick Willems, Jared T. Simpson, Mark D. Shriver, Jeffrey A. Rosenfeld, Carlos D. Bustamante, Stephen B. Montgomery, Francisco M. De La Vega, Jake K. Byrnes, Andrew W. Carroll, Marianne K. DeGorter, Phil Lacroute, Brian K. Maples, Alicia R. Martin, Andres Moreno-Estrada, Suyash S. Shringarpure, Fouad Zakharia, Eran Halperin, Yael Baran, Charles Lee, Eliza Cerveira, Jaeho Hwang, Ankit Malhotra, Dariusz Plewczynski, Kamen Radew, Mallory Romanovitch, Chengsheng Zhang, Fiona C. L. Hyland, David W. Craig, Alexis Christoforides, Nils Homer, Tyler Izatt, Ahmet A. Kurdoglu, Shripad A. Sinari, Kevin Squire, Stephen T. Sherry, Chunlin Xiao, Jonathan Sebat, Danny Antaki, Madhusudan Gujral, Amina Noor, Kenny Ye, Esteban G. Burchard, Ryan D. Hernandez, Christopher R. Gignoux, David Haussler, Sol J. Katzman, W. James Kent, Bryan Howie, Andres Ruiz-Linares, Emmanouil T. Dermitzakis, Scott E. Devine, Gonçalo R. Abecasis, Hyun Min Kang, Jeffrey M. Kidd, Tom Blackwell, Sean Caron, Wei Chen, Sarah Emery, Lars Fritsche, Christian Fuchsberger, Goo Jun, Bingshan Li, Robert Lyons, Chris Scheller, Carlo Sidore, Shiya Song, Elzbieta Sliwerska, Daniel Taliun, Adrian Tan, Ryan Welch, Mary Kate Wing, Xiaowei Zhan, Philip Awadalla, Alan Hodgkinson, Yun Li, Xinghua Shi, Andrew Quitadamo, Gert. A global reference for human genetic variation. Nature, 526(7571):68–74, oct 2015.

[21] Michael A Gimbrone, Guillermo García-Cardeña, and Guillermo García-Cardeña. Endothelial Cell Dysfunction and the Pathobiology of Atherosclerosis. Circulation research, 118(4):620–36, feb 2016.

[22] Alexander Gusev, Arthur Ko, Huwenbo Shi, Gaurav Bhatia, Wonil Chung, Brenda W J H Penninx, Rick Jansen, Eco J C de Geus, Dorret I Boomsma, Fred A Wright, Patrick F Sullivan, Elina Nikkola, Marcus Alvarez, Mete Civelek, Aldons J Lusis, Terho Lehtimäki, Emma Raitoharju, Mika Kähönen, Ilkka Seppälä, Olli T Raitakari, Johanna Kuusisto, Markku Laakso, Alkes L Price, Päivi Pajukanta, and Bogdan Pasaniuc. Integrative approaches for large-scale transcriptome-wide association studies. Nature Genetics, 48(3):245–252, mar 2016.

[23] Hadi A R Hadi, Cornelia S Carr, and Jassim Al Suwaidi. Endothelial dysfunction: cardiovascular risk factors, therapy, and outcome. Vascular health and risk management, 1(3):183–98, 2005.

[24] Takashi Hirose and H Robert Horvitz. An Sp1 transcription factor coordinates caspase-dependent and -independent apoptotic pathways. Nature, 500(7462):354–8, aug 2013.

[25] Farhad Hormozdiari, Martijn van de Bunt, Ayellet V. Segre, Xiao Li, Jong Wha J. Joo, Michael Bilow, Jae Hoon Sul, Sriram Sankararaman, Bogdan Pasaniuc, and Eleazar Eskin. Colocalization of GWAS and eQTL Signals Detects Target Genes. The American Journal of Human Genetics, 99(6):1245–1260, dec 2016.

[26] Daehwan Kim, Ben Langmead, and Steven L Salzberg. HISAT: a fast spliced aligner with low memory requirements. Nature methods, 12(4):357–60, apr 2015.

[27] David A Knowles, Courtney K Burrows, John D Blischak, Kristen M Patterson, Daniel J Serie, Nadine Norton, Carole Ober, Jonathan K Pritchard, and Yoav Gilad. Determining the genetic basis of anthracycline-cardiotoxicity by molecular response QTL mapping in induced cardiomyocytes. eLife, 7, may 2018.

[28] David A Knowles, Joe R Davis, Hilary Edgington, Anil Raj, Marie-Julie Favé, Xiaowei Zhu, James B Potash, Myrna M Weissman, Jianxin Shi, Douglas F Levinson, Philip Awadalla, Sara Mostafavi, Stephen B Montgomery, and Alexis Battle. Allele-specific expression reveals interactions between genetic variation and environment. Nat. Methods, 14(7):699–702, may 2017.

[29] Hua Li, Sheng-Yu Jin, Hyun-Joon Son, Je Hoon Seo, and Goo-Bo Jeong. Caffeine-induced endothelial cell death and the inhibition of angiogenesis. Anatomy & cell biology, 46(1):57–67, mar 2013.

[30] Michael I Love, Wolfgang Huber, and Simon Anders. Moderated estimation of fold change and dispersion for RNA-seq data with DESeq2. Genome biology, 15(12):550, dec 2014.

[31] Xin M Luo and A Catharine Ross. Retinoic acid exerts dual regulatory actions on the expression and nuclear localization of interferon regulatory factor-1. Experimental biology and medicine (Maywood, N.J.), 231(5):619–31, may 2006.

[32] Shozo Matsuoka, Toshitake Moriyama, Noriyuki Ohara, Kenji Tanimura, and Takeshi Maruo. Caffeine induces apoptosis of human umbilical vein endothelial cells through the caspase-9 pathway. Gynecological endocrinology : the official journal of the International Society of Gynecological Endocrinology, 22(1):48–53, jan 2006.

[33] G A Moyerbrailean, G O Davis, C T Harvey, D Watza, X Wen, R Pique-Regi, and F Luca. A high-throughput RNA-seq approach to profile transcriptional responses. Sci. Rep., 5:14976, oct 2015.

[34] Gregory A. Moyerbrailean, Cynthia A. Kalita, Chris T. Harvey, Xiaoquan Wen, Francesca Luca, and Roger Pique-Regi. Which Genetics Variants in DNase-Seq Footprints Are More Likely to Alter Binding? PLOS Genetics, 12(2):e1005875, feb 2016.

[35] Gregory A. Moyerbrailean, Allison L. Richards, Daniel Kurtz, Cynthia A. Kalita, Gordon O. Davis, Chris T. Harvey, Adnan Alazizi, Donovan Watza, Yoram Sorokin, Nancy Hauff, Xiang Zhou, Xiaoquan Wen, Roger Pique-Regi, and Francesca Luca. High-throughput allele-specific expression across 250 environmental conditions. Genome Research, 26(12):1627–1638, dec 2016.

[36] Yohann Nédélec, Joaquín Sanz, Golshid Baharian, Zachary A. Szpiech, Alain Pacis, Anne Dumaine, Jean-Christophe Grenier, Andrew Freiman, Aaron J. Sams, Steven Hebert, Ariane Pagé Sabourin, Francesca Luca, Ran Blekhman, Ryan D. Hernandez, Roger Pique-Regi, Jenny Tung, Vania Yotova, and Luis B. Barreiro. Genetic Ancestry and Natural Selection Drive Population Differences in Immune Responses to Pathogens. Cell, 167(3):657–669.e21, oct 2016.

[37] Majid Nikpay, Anuj Goel, Hong-Hee Won, Leanne M Hall, Christina Willenborg, Stavroula Kanoni, Danish Saleheen, Theodosios Kyriakou, Christopher P Nelson, Jemma C Hopewell, Thomas R Webb, Lingyao Zeng, Abbas Dehghan, Maris Alver, Sebastian M Armasu, Kirsi Auro, Andrew Bjonnes, Daniel I Chasman, Shufeng Chen, Ian Ford, Nora Franceschini, Christian Gieger, Christopher Grace, Stefan Gustafsson, Jie Huang, Shih-Jen Hwang, Yun Kyoung Kim, Marcus E Kleber, King Wai Lau, Xiangfeng Lu, Yingchang Lu, Leo-Pekka Lyytikäinen, Evelin Mihailov, Alanna C Morrison, Natalia Pervjakova, Liming Qu, Lynda M Rose, Elias Salfati, Richa Saxena, Markus Scholz, Albert V Smith, Emmi Tikkanen, Andre Uitterlinden, Xueli Yang, Weihua Zhang, Wei Zhao, Mariza de Andrade, Paul S de Vries, Natalie R van Zuydam, Sonia S Anand, Lars Bertram, Frank Beutner, George Dedoussis, Philippe Frossard, Dominique Gauguier, Alison H Goodall, Omri Gottesman, Marc Haber, Bok-Ghee Han, Jianfeng Huang, Shapour Jalilzadeh, Thorsten Kessler, Inke R König, Lars Lannfelt, Wolfgang Lieb, Lars Lind, Cecilia M Lindgren, Marja-Liisa Lokki, Patrik K Magnusson, Nadeem H Mallick, Narinder Mehra, Thomas Meitinger, Fazal-ur-Rehman Memon, Andrew P Morris, Markku S Nieminen, Nancy L Pedersen, Annette Peters, Loukianos S Rallidis, Asif Rasheed, Maria Samuel, Svati H Shah, Juha Sinisalo, Kathleen E Stirrups, Stella Trompet, Laiyuan Wang, Khan S Zaman, Diego Ardissino, Eric Boerwinkle, Ingrid B Borecki, Erwin P Bottinger, Julie E Buring, John C Chambers, Rory Collins, L Adrienne Cupples, John Danesh, Ilja Demuth, Roberto Elosua, Stephen E Epstein, Tõnu Esko, Mary F Feitosa, Oscar H Franco, Maria Grazia Franzosi, Christopher B Granger, Dongfeng Gu, Vilmundur Gudnason, Alistair S Hall, Anders Hamsten, Tamara B Harris, Stanley L Hazen, Christian Hengstenberg, Albert Hofman, Erik Ingelsson, Carlos Iribarren, J Wouter Jukema, Pekka J Karhunen, Bong-Jo Kim, Jaspal S Kooner, Iftikhar J Kullo, Terho Lehtimäki, Ruth J F Loos, Olle Melander, Andres Metspalu, Winfried März, Colin N Palmer, Markus Perola, Thomas Quertermous, Daniel J Rader, Paul M Ridker, Samuli Ripatti, Robert Roberts, Veikko Salomaa, Dharambir K Sanghera, Stephen M Schwartz, Udo Seedorf, Alexandre F Stewart, David J Stott, Joachim Thiery, Pierre A Zalloua, Christopher J O’Donnell, Muredach P Reilly, Themistocles L Assimes, John R Thompson, Jeanette Erdmann, Robert Clarke, Hugh Watkins, Sekar Kathiresan, Ruth McPherson, Panos Deloukas, Heribert Schunkert, Nilesh J Samani, and Martin Farrall. A comprehensive 1000) Genomes?based genome-wide association meta-analysis of coronary artery disease. Nature Genetics, 47(10):1121–1130, oct 2015.

[38] Robert H. Oakley and John A. Cidlowski. The biology of the glucocorticoid receptor: New signaling mechanisms in health and disease. Journal of Allergy and Clinical Immunology, 132(5):1033–1044, nov 2013.

[39] Alain Pacis, Ludovic Tailleux, Alexander M Morin, John Lambourne, Julia L MacIsaac, Vania Yotova, Anne Dumaine, Anne Danckaert, Francesca Luca, Jean-Christophe Grenier, Kasper D Hansen, Brigitte Gicquel, Miao Yu, Athma Pai, Chuan He, Jenny Tung, Tomi Pastinen, Michael S Kobor, Roger Pique-Regi, Yoav Gilad, and Luis B Barreiro. Bacterial infection remodels the DNA methylation landscape of human dendritic cells. Genome Res., 25(12):1801–11, dec 2015.

[40] Jing Pan and Kenneth M. Baker. Retinoic Acid and the Heart. In Vitamins and hormones, volume 75, pages 257–283. 2007.

[41] Roger Pique-Regi, Jacob F Degner, Athma a Pai, Daniel J Gaffney, Yoav Gilad, and Jonathan K Pritchard. Accurate inference of transcription factor binding from DNA sequence and chromatin accessibility data. Genome Res., 21(3):447–55, mar 2011.

[42] Peramaiyan Rajendran, Thamaraiselvan Rengarajan, Jayakumar Thangavel, Yutaka Nishigaki, Dhanapal Sakthisekaran, Gautam Sethi, and Ikuo Nishigaki. The Vascular Endothelium and Human Diseases. International Journal of Biological Sciences, 9(10):1057–1069, 2013.

[43] Margaret P Rayman. Selenium and human health. Lancet (London, England), 379(9822):1256–68, mar 2012.

[44] Marco Roman, Petru Jitaru, and Carlo Barbante. Selenium biochemistry and its role for human health. Metallomics : integrated biometal science, 6(1):25–54, jan 2014.

[45] Akiko Saito, Akira Sugawara, Akira Uruno, Masataka Kudo, Hiroyuki Kagechika, Yasufumi Sato, Yuji Owada, Hisatake Kondo, Mayumi Sato, Masahiko Kurabayashi, Masue Imaizumi, Shigeru Tsuchiya, and Sadayoshi Ito. All-trans Retinoic Acid Induces in Vitro Angiogenesis via Retinoic Acid Receptor: Possible Involvement of Paracrine Effects of Endogenous Vascular Endothelial Growth Factor Signaling. Endocrinology, 148(3):1412–1423, mar 2007.

[46] Robert M. Sapolsky, L. Michael Romero, and Allan U. Munck. How Do Glucocorticoids Influence Stress Responses? Integrating Permissive, Suppressive, Stimulatory, and Preparative Actions. Endocrine Reviews, 21(1):55–89, feb 2000.

[47] E Schoenmakers, G Verrijdt, B Peeters, G Verhoeven, W Rombauts, and F Claessens. Differences in DNA binding characteristics of the androgen and glucocorticoid receptors can determine hormone-specific responses. The Journal of biological chemistry, 275(16):12290–7, apr 2000.

[48] Tesa M. Severson, Yongsoo Kim, Stacey E. P. Joosten, Karianne Schuurman, Petra van der Groep, Cathy B. Moelans, Natalie D. ter Hoeve, Quirine F. Manson, John W. Martens, Carolien H. M. van Deurzen, Ellis Barbe, Ingrid Hedenfalk, Peter Bult, Vincent T. H. B. M. Smit, Sabine C. Linn, Paul J. van Diest, Lodewyk Wessels, and Wilbert Zwart. Characterizing steroid hormone receptor chromatin binding landscapes in male and female breast cancer. Nature Communications, 9(1):482, dec 2018.

[49] Barbara E Stranger, Alexandra C Nica, Matthew S Forrest, Antigone Dimas, Christine P Bird, Claude Beazley, Catherine E Ingle, Mark Dunning, Paul Flicek, Daphne Koller, Stephen Montgomery, Simon Tavaré, Panos Deloukas, and Emmanouil T Dermitzakis. Population genomics of human gene expression. Nat. Genet., 39(10):1217–24, oct 2007.

[50] Duncan Turnbull, Joseph V. Rodricks, Gregory F. Mariano, and Farah Chowdhury. Caffeine and cardiovascular health. Regulatory Toxicology and Pharmacology, 89:165–185, oct 2017.

[51] Duncan Turnbull, Joseph V. Rodricks, Gregory F. Mariano, and Farah Chowdhury. Caffeine and cardiovascular health. Regulatory Toxicology and Pharmacology, 89:165–185, oct 2017.

[52] Brian R Walker. Glucocorticoids and Cardiovascular Disease. European Journal of Endocrinology, 157(5):545–559, nov 2007.

[53] Xiaoquan Wen, Yeji Lee, Francesca Luca, and Roger Pique-Regi. Efficient Integrative Multi-SNP Association Analysis via Deterministic Approximation of Posteriors. Am. J. Hum. Genet., 98(6):1114–1129, jun 2016.

[54] Xiaoquan Wen, Roger Pique-Regi, and Francesca Luca. Integrating molecular QTL data into genomewide genetic association analysis: Probabilistic assessment of enrichment and colocalization. PLOS Genetics, 13(3):e1006646, mar 2017.

[55] Daria V Zhernakova, Patrick Deelen, Martijn Vermaat, Maarten van Iterson, Michiel van Galen, Wibowo Arindrarto, Peter van ‘t Hof, Hailiang Mei, Freerk van Dijk, Harm-Jan Westra, Marc Jan Bonder, Jeroen van Rooij, Marijn Verkerk, P Mila Jhamai, Matthijs Moed, Szymon M Kielbasa, Jan Bot, Irene Nooren, René Pool, Jenny van Dongen, Jouke J Hottenga, Coen D A Stehouwer, Carla J H van der Kallen, Casper G Schalkwijk, Alexandra Zhernakova, Yang Li, Ettje F Tigchelaar, Niek de Klein, Marian Beekman, Joris Deelen, Diana van Heemst, Leonard H van den Berg, Albert Hofman, André G Uitterlinden, Marleen M J van Greevenbroek, Jan H Veldink, Dorret I Boomsma, Cornelia M van Duijn, Cisca Wijmenga, P Eline Slagboom, Morris A Swertz, Aaron Isaacs, Joyce B J van Meurs, Rick Jansen, Bastiaan T Heijmans, Peter A C ‘t Hoen, and Lude Franke. Identification of context-dependent expression quantitative trait loci in whole blood. Nat. Genet., 49(1):139–145, jan 2017.

[56] Dongxing Zhu, Nabil A. Rashdan, Karen E. Chapman, Patrick WF Hadoke, and Vicky E. MacRae. A novel role for the mineralocorticoid receptor in glucocorticoid driven vascular calcification. Vascular Pharmacology, 86:87–93, nov 2016.

